# Lack of robust evidence for a *Wolbachia* infection in *Anopheles gambiae* from Burkina Faso

**DOI:** 10.1101/2022.03.05.483130

**Authors:** Simon P. Sawadogo, Didier A. Kabore, Ezechiel B. Tibiri, Angela Hughes, Olivier Gnankine, Shannon Quek, Abdoulaye Diabaté, Hilary Ranson, Grant L. Hughes, Roch K. Dabiré

## Abstract

The endosymbiont *Wolbachia* can have major effects on the reproductive fitness, and vectorial capacity of host insects and may provide new avenues to control mosquito borne pathogens. *Anopheles gambiae* s.l is the major vector of malaria in Africa but the use of *Wolbachia* in this species has been limited by challenges in establishing stable transinfected lines and uncertainty around native infections. High frequencies of infection of *Wolbachia* have been previously reported in *An. gambiae* collected from the Valle du Kou region of Burkina Faso in 2011 and 2014. Here we re-evaluated the occurrence of *Wolbachia* in natural samples, collected from Valle du Kou over a 12-year time span, and in addition, expanded sampling to other sites in Burkina Faso. Our results showed that, in contrast to earlier reports, *Wolbachia* is present at an extremely low prevalence in natural population of *An. gambiae*. From 5,341 samples analysed only 29 were positive for *Wolbachia* by nested PCR representing 0.54% of prevalence. No positive samples were found with regular PCR. Phylogenetic analysis of 16S rRNA gene amplicons clustered across supergroup B, with some having similarity to sequences previously found in *Anopheles* from Burkina Faso. However, we cannot discount the possibility that the amplicon positive samples we detected were due to environmental contamination or were false positives. Regardless, the lack of a prominent native infection in *An. gambiae* s.l. is encouraging for applications utilising *Wolbachia* traninsfected mosquitoes for malaria control.

## Background

*Wolbachia* is an obligate intracellular bacterial symbiont, found in many insect species that, in recent years, has shown great potential use for use in vector-borne pathogen control. It is well known for its ability to manipulate host reproduction enabling it to spread into insect populations (Werren et al., 2008). Additionally, *Wolbachia* can inhibit the development of diverse pathogens (Hedges et al., 2008; Kambris et al., 2009; Moreira et al., 2009; Hughes et al., 2011; Kambris et al., 2010; Shaw et al., 2016) which makes *Wolbachia* an attractive agent for pathogen control. Some strains of *Wolbachia* protect insect hosts from viral infections (Chrostek et al., 2013; Hedges et al., 2008; Glaser et al., 2010) and the presence of *Wolbachia* in *Aedes aegypti* mosquitoes impair infections with dengue and other arboviruses (Walker et al., 2011; Moreira et al., 2009). Based on these findings, releases of *Wolbachia* infected male and female mosquitoes have been undertaken with the aim of spreading *Wolbachia*-mediated resistance to viruses in natural mosquito populations (Hoffmann et al., 2011) with initial clinical trials showing positive outcomes (Indriani et al., 2020; Utarini et al 2021). In addition to population replacement approaches, *Wolbachia* has also been exploited for population suppression control strategies whereby infected male are released to reduce mosquito numbers by inducing cytoplasmic incompatibility when mating with uninfected females (LAVEN, 1967; Zheng et al., 2019; Crawford et al., 2020)

In *Anopheles* mosquitoes there are several reports indicating amplification of the *Wolbachia* 16S rRNA gene fragment by nested PCR. In general, these studies find *Wolbachia* PCR positive individuals at low frequency and infection prevalence in the population (Baldini et al., 2014; Shaw et al 2016; Ayala et al., 2019; Gomez et al., 2017), however some studies reporting a negative correlation between amplicon positive mosquitoes and *Plasmodium* development in natural populations (Shaw et al., 2016; Gomez et al., 2017). Recently it has been shown that *Anopheles moucheti* and *An. demeilloni* possess high density infections with *Wolbachia* infections being observed in the germline, and complete genomes recovered (Jeffries et al., 2018; Walker et al., 2021; Quek et al., 2021). Significant efforts to establish artificially transinfected lines of *Anopheles* with *Wolbachia* have proved largely unsuccessful (Walker et al., 2011). Importantly, a stable line was established in *An. stephensi*, a vector of malaria in southern Asia, using *Wolbachia* from *Ae. albopictus* (*w*AlbB), which conferred resistance to *Plasmodium falciparum* infection (Bian et al., 2013). However, the *Wolbachia* infection induced fitness costs on the host, which would likely prevent establishment of the bacterium in mosquito populations (Joshi et al., 2014). Somatic, transient infections of the *Wolbachia* in *An. gambiae* were shown to significantly inhibit *P. falciparum* (Hughes et al., 2011), but the interference phenotype is variable (with other *Wolbachia* strain-parasite combinations (Hughes et al., 2012; 2014; Murdock et al., 2014). Interestingly, *Wolbachia* and other gut-associated microbes have negative associations (Hughes et al 2014; Rossi et al., 2015; Zink et al 2015), and these microbial interactions affect the biology of the host and transmission of the bacterium (Hughes et al., 2014), offering a possible reason for the lack of infection in some *Anopheles* species.

Despite the widespread report of amplification of *Wolbachia* by PCR in *Anopheles*, there is conjecture in the literature if *Anopheles* mosquitoes are truly infected with *Wolbachia* (Chrostek & Gerth, 2019). Most of the evidence stems from nested PCR approaches, a technique that is highly sensitive. This has led to suggestions that nested PCR may detect environmental *Wolbachia* DNA that the mosquito has encountered (Chrostek & Gerth, 2019). The low density and prevalence of infection and the phylogenetic diversity of strains reported indicates the infection is not stable in these associations or environmental DNA is being amplified. It is imperative to verify the prevalence of natural infections, and identify any native strains of *Wolbachia* in *Anopheles*, as these infections could impede population suppression or replacement control approaches exploiting transinfected lines. We previously reported high levels of natural infection in *An. gambiae* from Burkina Faso (Baldini et al., 2014; Shaw et al., 2016), but here we re-evaluate *An. gambiae* mosquitoes from these sites to examine the prevalence of infection in a wider geographical across a temporal sample set. Surprisingly we find low levels of amplicon positive mosquitoes, calling into question the findings from our earlier study.

## Methods

### Study area and mosquito collection

*Anopheles gambiae s.l*. mosquitoes were collected in seven villages during the rainy seasons of 2006, 2011, 2012, 2015, 2016 and 2018. These sites are located in the Sahelian zone (Kongoussi and Yilou) and Sudan-savanna zone (VK3, VK5, VK7, Soumousso and Tiefora) (Figure 1) of Burkina Faso. Adult mosquitoes were collected from the resting sites (inhabited houses, uninhabited houses, wood piles, clay pots) using mechanical aspirators or CDC light traps; males were also collected from mating swarms using an insect net (Diabaté et al., 2006). Adults were morphologically identified using the standard taxonomic key (Gillies & Coetzee, 1987, Gillies & de Meillon, 1968). Samples were stored at −20°C prior to molecular analyses.

**Figure 1:**
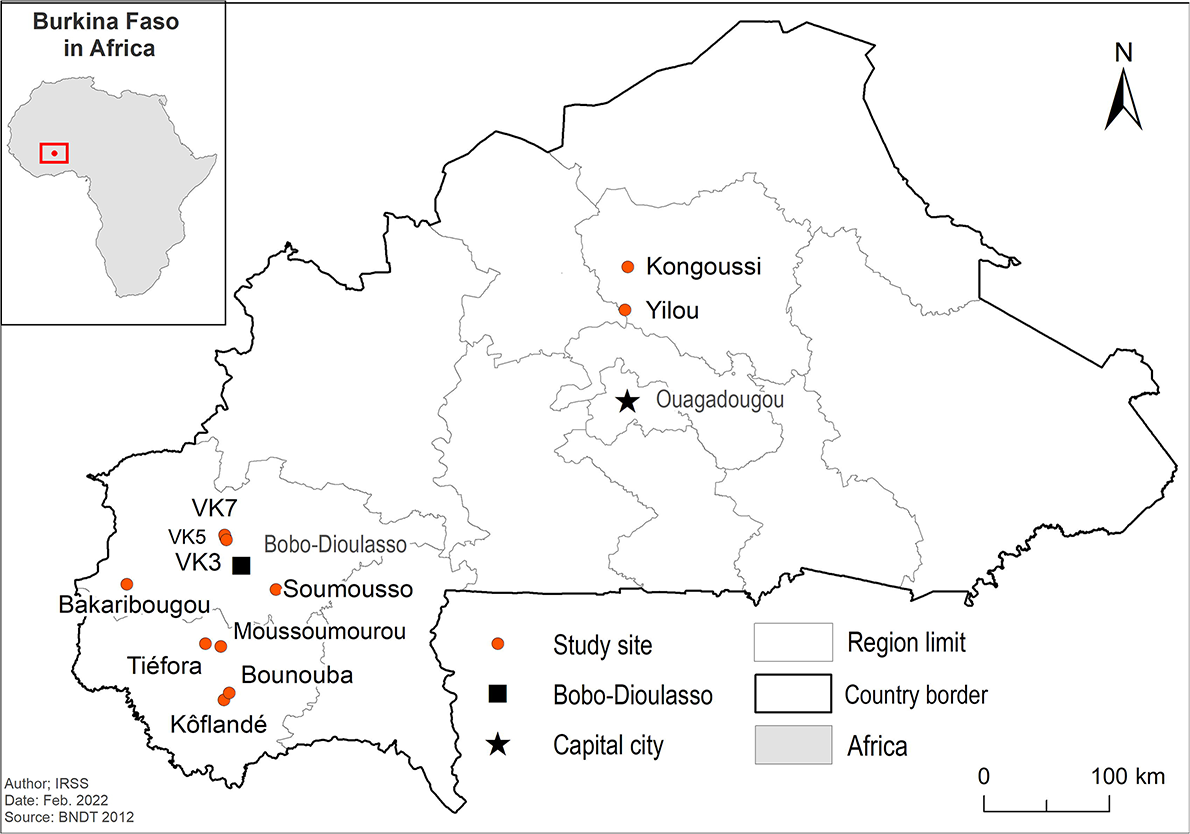
Map of Burkina Faso showing mosquito sampling sites.

### Mosquitoes DNA extraction and molecular analysis

DNA was extracted from 4953 *An gambiae* s.l. mosquitoes using the CTAB 2% extraction method. PCR was used to identify the species, as described previously (Santolamazza et al., 2008). To screen for *Wolbachia,* conventional (Werren & Windsor, 2000) or nested-PCR using *Wolbachia* specific primers was used to amplify the 16S rRNA gene (Shaw et al., 2016). Amplicon positive samples were repeated to confirm infection and the PCR products purified by filtration using NucleoFast ®96 PCR DNA purification plate. For Sanger sequencing Positive (DNA from *Culex* mosquitoes collected in the city of Bobo-Dioulasso) and negative controls were included to rule out PCR contamination. Samples were sequenced by GenoScreen.

In parallel to the 4953 samples we analyzed, a total of 388 samples, all collected as larvae either from VK7 (n=268) or from the Cascades district (n=120) were analysed at LSTM. These Samples were collected between October 2011 and September 2014 and included 292 samples that had been exposed to deltamethrin (199 survivors, 93 dead) (see Supplementary file S1). All were tested using the 16S-Wspecf/Wspecr primers (Werren et al., 2000).

### Phylogenetic analysis

Phylogenetic analyses of *Wolbachia* samples were performed on the conserved 16S rRNA sequences. Accession numbers of the sequences obtained in the present study and used to construct the tree are listed in Table 1. Contigs obtained were cleaned and assembled *de novo* using Geneious v. 8.1.7 (Biomatters Ltd). All sequences were subjected to BLAST search tools in NCBI using Geneious and subsequently to pairwise sequence comparison (Altschup et al., 1990; Bao et al., 2014). The homologous sequences were retrieved for phylogenetic analysis based on BLAST results.

**Table 1:** Spatial and temporal variation of *Wolbachia* infection prevalence in Burkina Faso

| Year | Village | Species | Tested | Amplicons negative | Amplicons positive (First nested PCR) | Amplicons positive (repeated Nested PCR) |
| --- | --- | --- | --- | --- | --- | --- |
| 2006 | Soumouosso | *An. gambiae* | 331 | 310 | 24 (7.25%) | 21 (6.34%) |
|  | VK7 | *An. coluzzii* | 176 | 170 | 19 (11.18%) | 6 (3.41%) |
| 2011 | VK3 | *An. coluzzii* | 506 | 506 | 0 | 0 |
| 2015 | Soumouosso | *An. gambiae* | 20 | 20 | 0 | 0 |
| 2016 | Soumouosso | *An. gambiae* | 562 | 562 | 0 | 0 |
|  | VK5 | *An. coluzzii* | 868 | 867 | 1 (0.16%) | 0 |
| 2018 | Kongoussi | *An. coluzzii* | 3 | 3 | 0 | 0 |
|  | Soumouosso | *An. gambiae* | 237 | 237 | 0 | 0 |
|  | VK3 | *An. coluzzii* | 132 | 130 | 2 (1.52%) | 2 (1.52%) |
|  | VK5 | *An. coluzzii* | 853 | 853 | 0 | 0 |
|  | VK7 | *An. coluzzii* | 1200 | 1200 | 0 | 0 |
|  | Yilou | *An. coluzzii* | 65 | 65 | 0 | 0 |
| **Total** |  |  | **4953** | **4923** | **46 (0.93%)** | **29 (0.59%)** |

From this, we obtained a total of 70 sequences of 16S RNA from *Wolbachia* of supergroups A and B. This number also included 13 sequences that were reportedly isolated from *An. gambiae*. These 70 sequences were added to the 17 sequences obtained from this study, as well as an additional sequence from *Wolbachia* of *Culex* mosquitoes, sequenced as part of this study. All of these 88 sequences were then used as input into the program MAFFT (v7.455 Katoh and Standley, 2013), and aligned using default parameters. The resultant alignment was then manually processed to remove ambiguously aligned regions at the 5’ and 3’ ends, as well as any columns that contained undetermined nucleotides from sequencing. This curated alignment was then used as input into the program IQTree (v1.6.1, Nguyen et al., 2015) to build a phylogenetic tree, utilising the DNA substitution model K2P+G4 (best-fit model as determined by the ModelFinder algorithm, Kalyaanamoorthy et al., 2017), with 1,000 non-parametric bootstrap replicates. The tree was then visualized and edited using FigTree v.1.4.4.

## Results

No amplicon positive samples were found with the regular PCR but a total of 46 of the 4,953 amplicons were positive for *Wolbachia* using the more sensitive nested PCR representing a prevalence of 0.93%. The nested PCR was repeated for each of the initial 46 positive mosquitoes and 29 remained positive on the repeat PCR, representing a prevalence of 0.59% (29/4953). The inability to confirm 17 individuals in a repeated nested-PCR suggested the template is at the limits of detection in these samples.

*Wolbachia* was only detected in samples from two of the six years; in 2006, *Wolbachia* prevalence was 5.33 % (n= 507) and 0.08 % (n=2490) in 2018 (Table 1). Regarding the spatial distribution, all positive specimens were found in just three of the seven villages (VK3, VK7 and Soumoussso), all located in the Sudan savanna zone in the western region of Burkina Faso. The prevalence of positive samples was low with 0.31% (2/638) in VK3, 0.36% (6/1376) in VK7 and 1.87%: (21/1150) in Soumousso. A significantly higher proportion of *An. gambiae* s.s were amplicon positive for *Wolbachia* than *An. coluzzii* (χ^2^ = 23.493, df = 1, p < 0.0001; *An*. *gambiae* 2.08% (24/1150); *An. coluzzii* 0.53% (22/4151)

Concerning the 388 samples analyzed in LSTM from VK7 and Tiefora no positive amplicon was found in any sample. However, positives bands were observed in positive control using the same protocol.

### Phylogenetic Analysis of *Wolbachia*

Phylogenetic analysis of the 16S rRNA gene sequences used in this study (70 published sequences, 17 from this study, one positive control from *Wolbachia* of *Culex pipiens*) resulted in a curated alignment of 317 nucleotides in length. A midpoint-rooted tree showed that all sequences obtained from this study clustered with supergroup B *Wolbachia*. From this study, 12 sequences of *An. gambiae* and *An. coluzzii* were noted to cluster into a weakly supported clade (Figure 2) alongside *w*AlbB, and six previously published sequences collected from *An. gambiae* in Burkina Faso and Guinea. The remaining six sequences obtained from this study were observed to be distributed throughout supergoup B, with none in supergoup A. Contrasting with the sequences from this study, previously published sequences from *An. gambiae* were observed to be distributed throughout both supergroups A and B (Figure 2).

**Figure 2:**
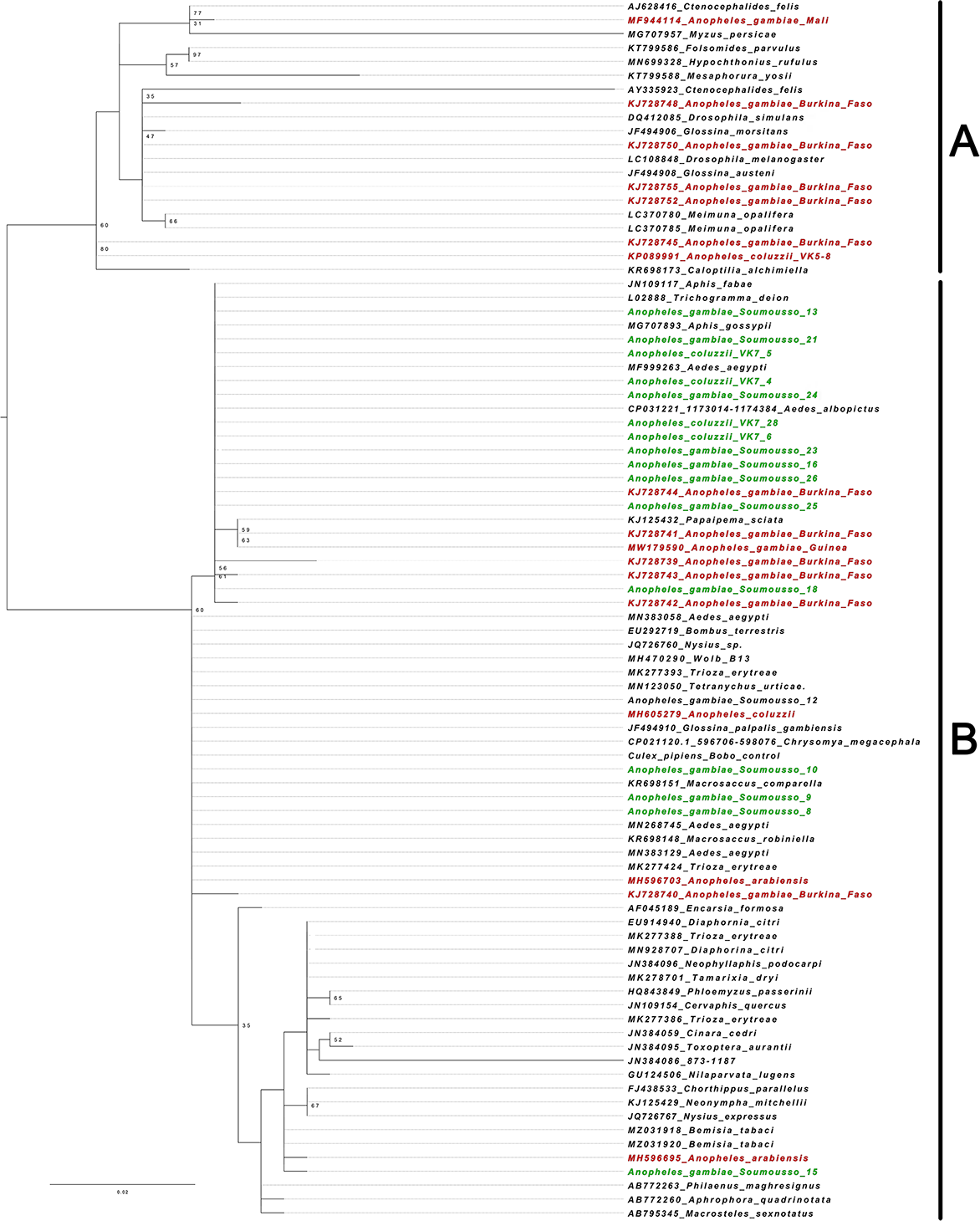
Phylogenetic Analysis of *Wolbachia* Strain. The green colour represents the sequences found in our samples, whilst red colour represents sequences previously identified in *Anopheles gambiae*.

When the alignment was reduced to look at the 18 sequences from this study, a total of six variant sites were observed, with five of these sites being unique to one sequence (Supplementary file S2, sample Sawadogo-15). Expanding this alignment to include all sequences from *Anopheles* mosquitoes, as well as *Wolbachia* from *A. albopictus*, a total of 31 variant sites were observed. The majority of these variant sites were observed to occur within seven of the published *Wolbachia* sequences, all of which cluster within supergroup A of the phylogenetic tree.

## Discussion

Despite being common among *Culex* and some *Aedes* mosquitoes (Carvajal et al., 2019), *Wolbachia* infections are relatively rare in most *Anopheles species*. While there are emerging reports suggesting that some *Anopheles* possess low prevalence infections (Baldini et al., 2014; Shaw et al., 2016; Ayala et al., 2019; Gomez et al., 2017), these studies use nested PCR which is a highly sensitive technique that could amplify environmental DNA. As such the veracity of these studies has been questioned (Chrostek and Gerth 2019). In contrast, a recent study demonstrated that *An. moucheti* and *An. demeilloni* possessed high-density maternally transmitted *Wolbachia* strains (Walker et al., 2021). Further genomic analysis of these strains indicated similar metabolic pathways to other *Wolbachia* strain and a close relationship to Wolbachia strains found in *Drosophila simulans* and *D. mauritiana* (Quek et al., 2022). Our expanded survey of *Wolbachia* from *An. gambiae* collected in Burkina Faso found an extremely low prevalence of *Wolbachia* using the nested-PCR screening approach.

The intensity of infection in most *Anopheles* is far lower than found in other insects. Analysis of 4,953 samples collected over a twelve-year period in Burkina Fasu found only 0.59% (29/4953) were amplicon positive for *Wolbachia*. This contrasts with an infection rate of 46% (275/602) in *An. coluzzii* in 2014 in the same area (Shaw et al., 2016) and varying spatially and temporally with samples from Mali with a minimal prevalence of 45% in Dangassa in 2015 reaching 95% (38/40) in *An. gambiae* sl in Kenieroba in 2016 (Gomez et al., 2017). A separate screening conducted in parallel at the Liverpool School of Tropical Medicine, to detect *Wolbachia* in *An. gambiae* sl. survivors of pyrethroid exposure collected in 2011 and 2012 from the same geographic region, failed to detect any *Wolbachia* samples in 348 individuals screened. Corroborating our findings, more recent studies using amplicon sequencing of the 16S rRNA gene have found minimal evidence for *Wolbachia* infection of *An. gambiae* in Burkina Faso. The relative abundance of the *Wolbachia* in the *An. gambiae* microbiome was 0.00002% (Zoure et al., 2019), while amplicon sequencing of nested-PCR positive individuals only found 42 *Wolbachia* reads constituting 0.04% relative abundance of the microbiome (Straub et al., 2020). Phylogenetic analysis indicated low levels of diversity within the 18 sequences obtained in this study, with sequences from both *An. gambiae* and *An. coluzzii* showing few variants. Furthermore, a total of 12 of these sequences appeared identical to that found from *w*AlbB. Interestingly, several sequences originally isolated from *An. gambiae* also appeared closely related to this group. One of the 18 sequences identified in this study did show significant divergence, clustering with sequences found within other insects.

While some of our amplicons were Sanger sequenced and had similarity to *Wolbachia* we cannot discount the possibility these were false positives. While we did not explicitly determine the specificity rate of this nested PCR, presuming it had a specificity rate of 99%, we would expect on average 49 false positive PCRs. These figures are in line with our 29 positive samples. Similarly, these amplicon positives samples, several of which appear to be on the limits of detection, could have been due to amplification of amplified environmental DNA encoding for the *wsp* gene. For example, it is known that microbial DNA can persist within soil for years (Nielsen et al., 2007). Finally, it is possible that the signal was due to *Wolbachia* bacteria that were transiently associated with the mosquito. This may be facilitated by ectoparasitic mites or midges, infection with endoparasitic nematodes, cohabitation with other arthropods hosting these bacteria, or acquired through the nectar feeding in plants (Chrostek et al 2017). Interestingly, while our current study shows that *Wolbachia* is present at a very low prevalence in *An. gambiae*, we saw more amplicon positive samples from villages (Soumousso (6.34%) and VK7 (3.41%)) where it had been detected previously in 2006 (Baldini et al., 2014; Shaw et al., 2016). This could possible indicate there was a biological factor, such as a *Wolbachia*-infected arthropod that cohabitated with *An. gambiae*, in this region and time producing amplicon positive signals. As low prevalence, low intensity infections are challenging to accurately detect it is not possible to conclude with certainty that the *Wolbachia* detected in this study, or our earlier published studies (Baldini et al., 2014; Shaw et al., 2016) are native infections of *Anopheles*.

## Author’s Contributions

SPS, GLH, and RKD designed the investigations; SPS coordinated the field collection; SPS DAK and AH performed laboratory analysis of samples, SPS, SQ, EBT and OG undertake the phylogenetic Analysis; SPS, GLH, HR and RKD wrote the manuscript with the inputs from DAK, EBT, AH, SQ, OG and AD. All authors read and approved the final manuscript.

## Acknowledgements

We are grateful to the populations of the study sites for their sincere cooperation during mosquito sampling. This work was supported by the Partnership for Increasing the Impact of Vector Control (PIIVeC) funded by the Medical Research Council of the UK (grant number MR/P027873/1) through the Global Challenges Research Fund.

## Table

Supplementary Table 1: Details of mosquito samples from Burkina Faso tested for the presence of Wolbachia at LSTM

## Figures

Supplementary Figure 1: Single-nucleotide variant alignment of all sequences from *Anopheles gambiae*, and *Wolbachia* of *Aedes albopictus*.

## References

Altschup, S.F., Gish, W., Miller, W., Myers, E.W., & Lipman, D.J. (1990) Basic local alignment search tool. Journal of Molecular Biology, 215, 403–410

Ayala, D., Akone-EllA, O., Rahola, N., Kengne, P., Ngangue, M.F., Mezeme, F., Makanga, B.K., Nigg, M., Costantini, C., Simard, F., Prugnolle., Roche, B., Duron, O., & Paupy, C. (2019) Natural *Wolbachia* infections are common in the major malaria vectors in Central Africa. Evolutionary Applications, 12, 1583–1594

Baldini, F., Segata, N., Pompon, J., Marcenac, P., Shaw, W.R., Dabiré, R.K., Diabaté, A., Levashina, E.A., & Catteruccia, F. (2014) Evidence of natural *Wolbachia* infections in field populations of *Anopheles gambiae*. Nature Communication, 5, 3985.

Bao, Y., Chetvernin, V., & Tatusova, T. (2014) Improvements to pairwise sequence comparison (PASC): a genome-based web tool for virus classification. Archives of Virology, 159, 3293–3304.

Bian, G., Joshi, D., Dong, Y., Lu, P., Zhou, G., Pan, X., Xu, Y., Dimopoulos, G., & Xi, Z. (2013) *Wolbachia* invades *Anopheles stephensi* populations and induces refractoriness to Plasmodium infection. Science, 340, 748–751.

Bourtzis, K., Dobson, S.L., Xi, Z., Rasgon, J.L., Calvitti, M., Moreira, L.A., Bossin, H.C., Moretti, R., L Baton, L.A., Hughes, G.L., Mavingui, P., & Gilles, J.R.L. (2014) Harnessing mosquito-*Wolbachia* symbiosis for vector and disease control. Acta Tropica, 132, S150–S163.

Carvajal, T.M., Hashimoto, K., Harnandika, R.K., Amalin, D.M., & Watanabe, K. (2019) Detection of *Wolbachia* in field-collected *Aedes aegypti* mosquitoes in metropolitan Manila, Philippines. Parasites Vectors, 12, 361.

Chang, J.M., Tommaso, P., & Notredame, C. (2014) TCS: a new multiple sequence alignment reliability measure to estimate alignment accuracy and improve phylogenetic tree reconstruction. Molecular Biology and Evolution, 31, 1625–1637.

Chrostek, E., & Gerth, M. (2019) Is *Anopheles gambiae* a natural host of *Wolbachia*? mBio, 10, e00784–19.

Chrostek, E., Marialva, M.S.P., Esteves, S.S., Weinert, L.A., Martinez, J., Jiggins, F.M., & Teixeira, L. (2013) *Wolbachia* variants induce differential protection to viruses in Drosophila melanogaster: a phenotypic and phylogenomic analysis. PLoS Genetics, 9, e1003896.

Chrostek, E., Pelz-Stelinski, K., Hurst, G.G.D., & Hughes, G.L. (2017) Horizontal transmission of intracellular insect symbionts via plants. Frontier Microbiology, 8, 2237.

Crawford, J.E., Clarke, D.W., Criswell. V., Mark Desnoyer, M., Devon Cornel, D., et al. (2020) Efficient production of male *Wolbachia*-infected Aedes aegypti mosquitoes enables large-scale suppression of wild populations. Nature Biotechnology, 38, 482–492.

Gillies, M.T., & Coetzee, M. (1987) A supplement to the Anophelinae of Africa South of the Sahara (Afrotropical Region). In Publications of the South African Institute of Medical Research, Johnenesburg, South Africa, 55, 143p.

Gillies, M.T., & de Meillon, B. (1968) The Anophelinae of Africa South of the Sahara (Ethiopan Zoogeographical Region). Publications of the South African Institute for Medical Research, 54, 343p.

Glaser, R.L., & Meola, M.A. (2010) The native Wolbachia endosymbionts of *Drosophila melanogaster* and *Culex quinquefasciatus* increase host resistance to West Nile virus infection. PLoS ONE, 5, e11977.

Gomes, F.M., Hixson, B.L., Tyner, M.D.W., Ramirez, J.L., Canepa, G.E., Alves E Silva, T.L., Molina-Cruz, A., Keita, M., Kane, F., Traoré, B., Sogoba, N., & Barillas-Mury C. (2017) Effect of naturally occurring Wolbachia in *Anopheles gambiae* s.l. mosquitoes from Mali on Plasmodium falciparum malaria transmission. PNAS, 114,12566–12571.

Hedges, L.M., Brownlie, J.C., O’Neill, S.L., & Johnson, K.N. (2008) *Wolbachia* and virus protection in insects. Science, 322, 702.

Hoffmann, A.A., Montgomery, B.L., Popovici, J., Iturbe-Ormaetxe, I., Johnson, P.H., Muzzi, F., Greenfield, M., Durkan, M., Leong, Y.S., Dong, Y., Cook, H., Axford, J., Callahan, A.G., Kenny, N., Omodei, C., McGraw, E.A., Ryan, P.A., Ritchie, S.A., Turelli, M., & O’Neill, S.L. (2011) Successful establishment of *Wolbachia* in *Aedes* populations to suppress dengue transmission. Nature, 476, 454–457

Hughes, G.L., Dodson, B.L., Johnson, R.M., Murdock, C.C., Tsujimoto, H., Suzuki, Y., Patt, A.A., Cui, L., Nossa, C.W., Barry, R.M., Sakamoto, J.M., Hornett, E.A., & Rasgon, J.L. (2014) Native microbiome impedes vertical transmission of *Wolbachia* in *Anopheles* mosquitoes. PNAS, 111, 12498–12503.

Hughes, G.L., Koga, R., Xue, P., Fukastu, T., & Rasgon, J.L. (2011) *Wolbachia* infections are virulent and inhibit the human malaria parasite *Plasmodium falciparum* in *Anopheles gambiae*. PLoS Pathogens, 7, e1002043.

Hughes, G.L., Vega-Rodriguez, J., Xue, P., & Rasgon, J.L. (2012) *Wolbachia* strain *w*AlbB enhances infection by the rodent malaria parasite *Plasmodium berghei* in *Anopheles gambiae* mosquitoes. Applied and Environmental Microbiology, 78, 1491–1495.

Indriani, C., Tantowijoyo, W., Rances, E., Andari, B., Prabowo, E., Yusdi, D., et al. (2020). Reduced dengue incidence following deployments of Wolbachia-infected Aedes aegypti in Yogyakarta, Indonesia: a quasi-experimental trial using controlled interrupted time series analysis. Gates Open Research, 4, 50.

Jeffries, C.L., Lawrence, G.G., Golovko, G., Kristan, M., Orsborne, J., Spence, K., Hurn, E., Bandibabone, J., Tantely, L.M., Raharimalala, F.N., Keita, K., Camara, D., Barry, Y., Wat’senga, F., Manzambi, E.Z., Afrane, Y.A., Mohammed, A.R., Abeku, T.A., Hedge, S., Khanipov, K., Pimenova, M., Fofanov, Y., Boyer, S., Irish, S.R., Hughes, G.L., & Walker, T. (2018) Novel *Wolbachia* strains in *Anopheles* malaria vectors from sub-Saharan Africa. Wellcome Open Research, 3, 113.

Jones, D.T., Taylor, W.R., & Thornton, J.M., (1992). The rapid generation of mutation data matrices from protein sequences. Computer Applications in the Biosciences, 8, 275–282.

Joshi, D., McFadden, M.J., Bevins, D., Zhang, F., & Xi, Z. (2014) *Wolbachia* strain *w*AlbB confers both fitness costs and benefit on *Anopheles stephensi*. Parasites Vectors, 7, 1–9.

Kalyaanamoorthy, S., Minh, B.Q., Wong, T.K.F., Haeseler, A., & Jermiin, L.S. (2017) ModelFinder: fast model selection for accurate phylogenetic estimates. Nature Methods, 14, 587–589.

Kambris, Z., Blagborough, A.M., Pinto, S.B., Blagrove, M.S., Godfray, H.C., Sinden, R.E., Sinkins, S.P. (2010) *Wolbachia* stimulates immune gene expression and inhibits plasmodium development in *Anopheles gambiae*. PLoS Pathogens, 6, e1001143.

Kambris, Z., Cook, P.E., Phuc, H.K., & Sinkins, S.P. (2009) Immune activation by life-shortening *Wolbachia* and reduced filarial competence in mosquitoes. Science, 326, 134–136.

Kumar, S., Stecher, G., Li, M., Knyaz, C., & Tamura, K. (2018) MEGA X: Molecular Evolutionary Genetics Analysis across computing platforms. Molecular Biology and Evolution, 35,1547–1549.

Kumar, S., Stecher, G., Peterson, D., & Tamura, K. (2012) MEGA-CC: Computing Core of Molecular Evolutionary Genetics Analysis Program for Automated and Iterative Data Analysis. Bioinformatics, 28, 2685–2686.

Laven, H. (1967) Eradication of *Culex pipiens fatigans* through Cytoplasmic Incompatibility. Nature, 216, 383–384.

Moreira, L.A., Iturbe-Ormaetxe, I., Jeffery, J.A., Lu, G., Pyke, A.T., Hedges. L.M., Rocha, B.C., Hall-Mendelin, S., Day, A., Riegler, M., Hugo, L.E., Johnson, K.N., Kay, B.H., McGraw, E.A., van den Hurk, A.F., Ryan, P.A., & O’Neill, S.L. (2009) A *Wolbachia* symbiont in *Aedes aegypti* limits infection with dengue, Chikungunya, and *Plasmodium*. Cell, 139, 1268–1278.

Murdock, C.C., Blanford, S., Hughes, G.L., Rasgon, J.L., & Thomas, M.B. (2014) Temperature alters *Plasmodium* blocking by *Wolbachia*. Scientific Reports, 4, 3932.

Niang, E.H.A., Bassene, H., Makoundou, P., Fenollar, F., Weill, M., & Mediannikov, O. (2018) First report of natural *Wolbachia* infection in wild *Anopheles funestus* population in Senegal. Malaria Journal, 17, 408.

Nielsen, K., Johnsen, P., Bensasson, D., & Daffonchio, D. (2007) Release and persistence of extracellular DNA in the environment. Environmental Biosafety Research, 6, 37–53.

Quek, S., Louise Cerdeira, L., Jeffries, C.L., Tomlinson, S., Walker, T., Hughes, G.L., & Heinz, E. (2021) *Wolbachia* endosymbionts in two *Anopheles* species indicates independent acquisitions and lack of prophage elements. bioRxiv, preprint doi: https://doi.org/10.1101/2021.11.15.468614; November 15, 2021.

Ranson, H., N’guessan, R., Lines, J., Moiroux, N., Nkuni, Z., & Corbel, V. (2011) Pyrethroid resistance in African anopheline mosquitoes: what are the implications for malaria control? Trends Parasitology, 27(2), 91–8.

Rasgon, J.L., Ren, X., & Petridis, M. (2006) Can *Anopheles gambiae* be infected with *Wolbachia pipientis*? Insights from an in vitro system. Applied and Environmental Microbiology, 72, 7718–7722.

Rossi, P., Ricci, I., Cappelli, A., Damiani, C., Ulissi, U. et al. (2015) Mutual exclusion of *Asaia* and *Wolbachia* in the reproductive organs of mosquito vectors. Parasites Vectors, 8, 278.

Santolamazza, F., Mancini, E., Simard, F., Qi, Y., Tu, Z., & Torre della, A. (2008) Insertion polymorphisms of SINE200 retro transposons within speciation islands of *Anopheles gambiae* molecular forms. Malaria journal, 7, 163.

Shaw, W.R., Marcenac, P., Childs, L.M., Buckee, C.O., Baldini, F., Sawadogo, S.P., Dabiré, R.K., Diabaté, A., & Catteruccia, F. (2016) *Wolbachia* infections in natural *Anopheles* populations affect egg laying and negatively correlate with *Plasmodium* development. Nature Communication, 7, 11772.

Straub, T.J., Shaw, W., Marcenac, P., Sawadogo., S.P., Dabire, R.K., Diabate, A., Catteruccia, F., & Neafsey, D.E. (2020) The *Anopheles coluzzii* microbiome and its interaction with the intracellular parasite *Wolbachia*. Scientific Reports, 10, 13847.

Walker, T., Quek, S., Jeffries, C.L., Bandibabone, J., Dhokiya, V.,et al. (2021) Stable high-density and maternally inherited Wolbachia infections in *Anopheles moucheti* and *Anopheles demeilloni* mosquitoes. Current Biology, 31, 1–11.

Utarini, A., Indriani, C., Ahmad, R.A., Tantowijoyo, W., Arguni, E., Ansari, M.R., Supriyati, E., Wardana, D.S., Meitika, Y., Ernesia, I., Nurhayati, I., Prabowo, E., Andari, B., Green, B.R., Hodgson, L., Cutcher, Z., Rancès, E., Ryan, P.A., O’Neill, S.L., Dufault, S.M., Tanamas, S.K., Jewell, N.P., Anders, K.L., & Simmons, C.P. (2021) Efficacy of *Wolbachia*-Infected Mosquito Deployments for the Control of Dengue. New England Journal of Medicine, 384, 2177–2186.

Walker, T., Johnson, P.H., Moreira, L.A., Iturbe-Ormaetxe, I, Frentiu, F.D., Mc-Meniman, C.J., Leong, Y.S., Dong, Y., Axford, J., Kriesner. P., Lloyd, A.L., Ritchie, S.A., O’Neill, S.L., & Hoffmann, A.A. (2011) The wMel *Wolbachia* strain blocks dengue and invades caged Aedes aegypti populations. Nature, 476, 450–453.

Werren, J.H., Baldo, L., & Clark, M.E. (2008) *Wolbachia*: master manipulators of invertebrate biology. Nature Reviews Microbiology, 6, 741–751.

Werren, J.H., & Windsor, D.M. (2000) *Wolbachia* infection frequencies in insects: evidence of a global equilibrium? Proceedings of the Royal Society B, 267, 1277–1285.

Zheng, X., Zhang, D., Li, Y., Yang, C., Wu, Y., et al. (2019) Incompatible and sterile insect techniques combined eliminate mosquitoes. Nature, 572, 56–61.

Zink, S.D., Van Slyke, G.A., Palumbo, M.J., Kramer, L.D., Ciota, A.T. (2015) Exposure to West Nile Virus Increases Bacterial Diversity and Immune Gene Expression in *Culex pipiens*. Viruses, 7(10), 5619–5631.

Zoure, A.A., Sare, A.R., Yameogo, F., Somda, Z., Massart, S., Badolo, A., & Francis, F. (2019) Bacterial communities associated with the midgut microbiota of wild *Anopheles gambiae* complex in Burkina Faso. Molecular Biology Reports, 47, 211–224.

