## Supplementary Table 1 for "Lack of robust evidence for a *Wolbachia* infection in *Anopheles gambiae* from Burkina Faso"

**Supplementary Table 1**: Details of mosquito samples from Burkina Faso tested for the presence of Wolbachia at LSTM

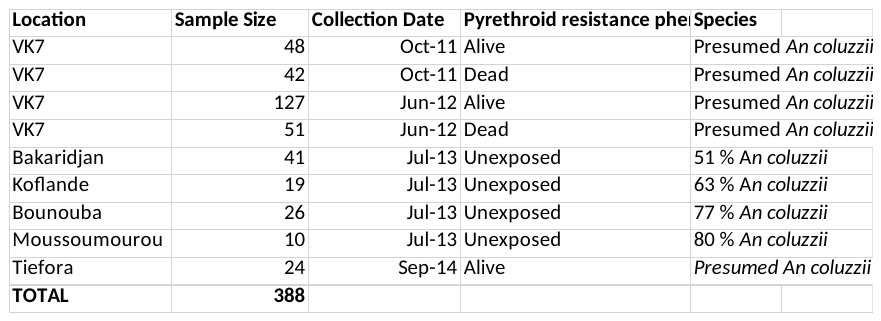
