## Supplementary Figure 1 for "Lack of robust evidence for a *Wolbachia* infection in *Anopheles gambiae* from Burkina Faso"

>ENA|AB772260/1024-1339

ttaagtcccgcaacgagcgcaaccctcatccttagttgcatcaggtaatgctgagcacttaaggaaactgcc

agtgataagctggaggaaggtggggatgatgtcaagtcatcatggcctttatgaagtgggctacacacgtgc

tacaatggtgtctacaatgggctgcaaggtgcgcaagcctaagctaat-ccctaaaagacatctcagttcgg

attgtactctgcaactcgagtgcatgaagttggaatcgctagtaatcgtggattagcatgccacggtgaata

cgttctcgggtcttgtacacactgcccgt

>ENA|AB772263/1024-1339

ttaagtcccgcaacgagcgcaaccctcatccttagttgcatcaggtaatgctgagcacttaaggaaactgcc

agtgataagctggaggaaggtggggatgatgtcaagtcatcatggcctttatgaagtgggctacacacgtgc

tacaatggtgtctacaatgggctgcaaggtgcgcaagcctaagctaat-ccctaaaagacatctcagttcgg

attgtactctgcaactcgagtgcatgaagttggaatcgctagtaatcgtggatcagcatgccacggtgaata

cgttctcgggtcttgtacacactgcccgt

>ENA|AB795345/1024-1339

ttaagtcccgcaacgagcgcaaccctcatccttagttgcatcaggtaaagctgagcacttaaggaaactgcc

agtgataagctggaggaaggtggggatgatgtcaagtcatcatggcctttatgaagtgggctacacacgtgc

tacaatggtgtctacaatgggctgcaaggtgcgcaagcctaagctaat-ccctaaaagacatctcagttcgg

attgtactctgcaactcgagtgcatgaagttggaatcgctagtaatcgtggatcagcatgccacggtgaata

cgttctcgggtcttgtacacactgcccgt

>ENA|AF045189/1033-1348

ttaagtcccgcaacgagcgcaaccctcttccttagttgsatcaggtaatgctgagtacttaaggaaactgcc

agtgataagctggaggaaggtggggatgatgtcaagtcatcatggsctttatgaagtgggctacacacgtgc

tacaatggtgtctacaatgggctgcaaggkgcgcaagcctaagctaat-ccctaaaagacatctcagttcgg

attgtactctgcaactcgagtgcatgaagttggaatcgctagtaatcgtggatcagcatgccacggtgaata

cgttctcgggtcttgtacacactgcccgt

>ENA|AJ628416/1021-1335

ttaagtcccgcaacgagcgcaaccctcatccttagttacatcaggtcatgctggggacttaaggaaactgcc

agtgataaactggaggaaggtggggatgatgtcaagtcatcatggcccttatggagtgggctacacacgtgc

tacaatggtggctacaatgggctgcaaagt-cgcaaggctgagctaat-ccttaaaagccatctcagttcgg

attgtactctgcaactcgagtgcatgaagttggaatcgctagtaatcgtggatcagcatgccacggtgaata

cgttctcgggtcttgtacacactgcccgt

>ENA|AY335923/960-1275

ttaagtcccgcaatgagggcaaccctcatccttagttacatcaggtaatgctggggacttaaggagactgcc

agtgatgaactggaggaaggtggggatgatgtcaagtcctcatgacccttacgggctgagctacacacgtgc

tacaatggtggctacaatgggctgcaaagt-cgcgaggctaagctaatcccttaaaagccatctcagttcgg

attgtactcagcaactcgagtgcatgaagtaggaatcgctagtaatcgtggatcagcacgccccggtgaata

cgttctcgggttttgtacacgctgaatgt

>ENA|DQ412085/999-1314

ttaagtcccgcaacgagcgcaaccctcatccttagttacatcaggtaatgctggggacttaaggaaactgcc

agtgataaactggaggaaggtggggatgatgtcaagtcatcatggcccttatggagtgggctacacacgtgc

tacaatggtggctacaatgggctgcaaagt-cgcgaggctaagctaatcccttaaaagccatctcagttcgg

attgtactctgcaactcgagtgcatgaagttggaatcgctagtaatcgtggatcagcacgccacggtgaata

cgttctcgggtcttgtacacactgcccgt

>ENA|EU292719/90-405

ttaagtcccgcaacgagcgcaaccctcatccttagttgcatcaggtaatgctgagtacttaaggaaactgcc

agtgataagctggaggaaggtggggatgatgtcaagtcatcatggcctttatggagtgggctacacacgtgc

tacaatggtgtctacaatgggctgcaaggtgcgcaagcctaagctaat-ccctaaaagacatctcagttcgg

attgtactctgcaactcgagtacatgaagttggaatcgctagtaatcgtggatcagcatgccacggtgaata

cgttctcgggtcttgtacacactgcccgt

>ENA|EU914940/593-908

ttaagtcccgcaacgagcgcaaccctcatccttagttgcatcaggtaatgctgagcacttaaggaaactgcc

agtgataagctggaggaagatggggatgatgtcaagtcatcatggcctttatgaagtgggctacacacgtgc

tacaatggtgtctacagtgggctgcaaggtgcgcaagcctaagctaat-ccctaaaagacatctcagttcgg

attgtactctgcaactcgagtgcatgaagttggaatcgctagtaatcgtggatcagcatgccacggtgaata

cgttctcgggtcttgtacacactgcccgt

>ENA|FJ438533/932-1247

ttaagtcccgcaacgagcgcaaccctcatccttagttgcatcaggtaatgctgagcacttaaggaaactccc

agtgataagctggaggaagatggggatgatgtcaagtcatcatggcctttatgaagtgggctacacacgtgc

tacaatggtgtctacaatgggctgcaaggtgcgcaagcctaagctaat-ccctaaaagacatctcagttcgg

attgtactctgcaactcgagtgcatgaagttggaatcgctagtaatcgtggatcagcatgccacggtgaata

cgttctcgggtcttgtacacactgcccgt

>ENA|GU124506/1044-1359

ttaagtcccgcaacgagcgcaaccctcatccttagttgcatcaggtaatgctgagcacttaaggaaactgcc

agtgataagctggaggaagatggggatgatgtcaagtcatcatggcctttatgaagtgggctacacacgtgc

tacaatggtgtctacagtgggctgcaaggtgcgcaagcctaagctaat-ccctaaaagacatctcagttcgg

attgtactctgcaactcgagtgcatgaagttggaatcgctagtaatcgtggatcagcatgccacggtgagta

cgttctcgggtcttgtacacactgcccgt

>ENA|HQ843849/782-1097

ttaagtcccgcaacgagcgcaaccctcatccttagttgcatcaggtaatgctgagcacttaaggaaactgcc

agtgataagctggaggaagatggggatgatgtcaagtcatcatggcctttatgaagtgggctacacacgtgc

tacaatggtgtctacagtgggctgcaaggtgcgcaagcctaggctaat-ccctaaaagacatctcagttcgg

attgtactctgcaactcgagtgcatgaagttggaatcgctagtaatcgtggatcagcatgccacggtgaata

cgttctcgggtcttgtacacactgcccgt

>ENA|JF494906/90-405

ttaagtcccgcaacgagcgcaaccctcatccttagttacatcaggtaatgctggggacttaaggaaactgcc

agtgataaactggaggaaggtggggatgatgtcaagtcatcatggcccttatggagtgggctacacacgtgc

tacaatggtggctacaatgggctgcaaagt-cgcgaggctaagctaatcccttaaaagccatctcagttcgg

attgtactctgcaactcgagtgcatgaagttggaatcgctagtaatcgtggatcagcacgccacggtgaata

cgttttcgggtcttgtacacactgcccgt

>ENA|JF494908/90-405

ttaagtcccgcaacgagcgcaaccctcatccttagttacatcaggtaatgctggggacttaaggaaactgcc

agtgataaactggaggaaggtggggatgatgtcaagtcatcatggcccttatggagtgggctacacacgtgc

tacaatggtggctacaatgggctgcaaagt-cgcgaggctaagctaatcccttaaaagccatctcagttcgg

attgtactctgcaactcgagtgcatgaagttggaatcgctagtaatcgtggatcagcacgccacggtgaata

cgttctcgggtcttgtacacactgcccgt

>ENA|JF494910/90-405

ttaagtcccgcaacgagcgcaaccctcatccttagttgcatcaggtaatgctgagtacttaaggaaactgcc

agtgataagctggaggaaggtggggatgatgtcaagtcatcatggcctttatggagtgggctacacacgtgc

tacaatggtgtctacaatgggctgcaaggtgcgcaagcctaagctaat-ccctaaaagacatctcagttcgg

attgtactctgcaactcgagtacatgaagttggaatcgctagtaatcgtggatcagcatgccacggtgaata

cgttctcgggtcttgtacacactgcccgt

>ENA|JN109117/90-405

ttaagtcccgcaacgagcgcaaccctcatccttagttgcatcaggtaatgctgagtacttaaggaaactgcc

agtgataagctggaggaaggtggggatgatgtcaagtcatcatggcctttatggagtgggctacacacgtgc

tacaatggtgtctacaatgggttgcaaggtgcgcaagcctaagctaat-ccctaaaagacatctcagttcgg

attgtactctgcaactcgagtacatgaagttggaatcgctagtaatcgtggatcagcatgccacggtgaata

cgttctcgggtcttgtacacactgcccgt

>ENA|JN109154/90-405

ttaagtcccgcaacgagcgcaaccctcatccttagttgcatcaggtaatgctgagcacttaaggaaactgcc

agtgataagctggaggaagatggggatgatgtcaagtcatcatggcctttatgaagtgggctacacacgtgc

tacaatggtgtctacagtgggctgcaaggtgcgcaagcctaggctaat-ccctaaaagacatctcagttcgg

attgtactctgcaactcgagtgcatgaagttggaatcgctagtaatcgtggatcagcatgccacggtgaata

cgttctcgggtcttgtacacactgcccgt

>ENA|JN384059/883-1198

ttaagtcccgcaacgagcgcaaccctcatccttagttgcatcaggtaatgctgagtacttaaggaaactgcc

agtgataagctggaggaagatggggatgatgtcaagtcatcatggcctttatgaagtgggctacacacgtgc

tacaatggtgtctacagtgggctgcaaggtgcgcaagcctaagctaat-ccctaaaagacatctcagttcgg

attgtactctgcaactcgagtgcatgaagttggaatcgctagtaatcgtggatcagcatgccacggtgaata

cgttctcgggtcttgtacacactgcccgt

>ENA|JN384086/873-1187

ttaagtcccgcaacgagcgcaaccctcatccttagttgcatcaggttatgctgaggacttaaggaaactgcc

agtgataagctggaggaagatggggatgatgtcaagtcatcatggcccttatgaagtgggctacacacgtgc

tacaatggtgcttacagtgggctgcaaggt-cgcaagcctaagctaat-cctaaaaaatcatctcagttcgg

attgttctctgcaactcgagagcatgaagttggaatcgctagtgatcgtggatcagcatgccacggtgaata

cgttctcgggtcttgtacacactgcccgt

>ENA|JN384095/691-1006

ttaagtcccgcaacgagcgcaaccctcatccttagttgcatcaggtaatgctgagtacttaaggaaactgcc

agtgataagctggaggaagatggggatgatgtcaagtcatcatggcctttatgaagtgggctacacacgtgc

tacaatggtgtctacagtaggctgcaaggtgcgcaagcctaagctaat-ccctaaaagacatctcagttcgg

attgtactctgcaactcgagtgcatgaagttggaatcgctagtaatcgtggatcagcatgccacggtgaata

cgttctcgggtcttgtacacactgcccgt

>ENA|JN384096/691-1006

ttaagtcccgcaacgagcgcaaccctcatccttagttgcatcaggtaatgctgagcacttaaggaaactgcc

agtgataagctggaggaagatggggatgatgtcaagtcatcatggcctttatgaagtgggctacacacgtgc

tacaatggtgtctacagtgggctgcaaggtgcgcaagcctaagctaat-ccctaaaagacatctcagttcgg

attgtactctgcaactcgagtgcatgaagttggaatcgctagtaatcgtggatcagcatgccacggtgaata

cgttctcgggtcttgtacacactgcccgt

>ENA|JQ726760/1025-1340

ttaagtcccgcaacgagcgcaaccctcatccttagttgcatcaggtaatgctgagtacttaaggaaactgcc

agtgataagctggaggaaggtggggatgatgtcaagtcatcatggcctttatggagtgggctacacacgtgc

tacaatggtgtctacaatgggctgcaaggtgcgcaagcctaagctaat-ccctaaaagacatctcagttcgg

attgtactctgcaactcgagtacatgaagttggaatcgctagtaatcgtggatcagcatgccacggtgaata

cgttctcgggtcttgtacacactgcccgt

>ENA|JQ726767/1024-1339

ttaagtcccgcaacgagcgcaaccctcatccttagttgcatcaggtaatgctgagcacttaaggaaactccc

agtgataagctggaggaagatggggatgatgtcaagtcatcatggcctttatgaagtgggctacacacgtgc

tacaatggtgtctacaatgggctgcaaggtgcgcaagcctaagctaat-ccctaaaagacatctcagttcgg

attgtactctgcaactcgagtgcatgaagttggaatcgctagtaatcgtggatcagcatgccacggtgaata

cgttctcgggtcttgtacacactgcccgt

>ENA|KJ125429/78-393

ttaagtcccgcaacgagcgcaaccctcatccttagttgcatcaggtaatgctgagcacttaaggaaactccc

agtgataagctggaggaagatggggatgatgtcaagtcatcatggcctttatgaagtgggctacacacgtgc

tacaatggtgtctacaatgggctgcaaggtgcgcaagcctaagctaat-ccctaaaagacatctcagttcgg

attgtactctgcaactcgagtgcatgaagttggaatcgctagtaatcgtggatcagcatgccacggtgaata

cgttctcgggtcttgtacacactgcccgt

>ENA|KJ125432/77-392

ttaagtcccgcaacgagcgcaaccctcatccttagttgcatcaggtaatgctgagtacttaaggaaactgcc

agtgataagctggaggaaggtggggatgatgtcaagtcatcatggcctttatggagtgggctacacacgtgc

tacaatggtgtctacaatgggttgcaaggtgcgcaagcytaagytaat-ccctaaaagacatctcagttcgg

attgtactctgcaactcgagtacatgaagttggaatcgctagtaatcgtggatcagcatgccacggtgaata

cgttctcgggtcttgtacacactgcccgt

>ENA|KJ728739/85-380

ttaagtcccgcaacgagcgcaaccctcatccttagttgcatcaggtaatgctgagtacttaaggaaactgcc

agtgataagctggaggaaggtggggatgatgtcaagtcatcatggcctttatggagtgggctacacacgtgc

tacaatggtgtctacaatgggttgcaaggtgcgcaagcctaagctaat-ccctaaaagacatctcagttcgg

attgtactctgcaactcgagtacatgaagttggaatcgctagtaatcgtggatcagcatgccacggtgaata

cgtctcggg--------------------

>ENA|KJ728740/91-406

ttaagtcccgcaacgagcgcaaccctcatccttagttgcatcaggtaatgctgaatacttaaggaaactgcc

agtgataagctggaggaaggtggggatgatgtcaagtcatcatggcctttatggagtgggctacacacgtgc

tacaatggtgtctacaatgggctgcaaggtgcgcaagcctaagctaat-ccctaaaagacatctcagttcgg

attgtactctgcaactcgagtacatgaagttggaatcgctagtaatcgtggatcaccatgccacggtgaata

cgttctcgggtcttgtacacactgcccgt

>ENA|KJ728741/91-394

ttaagtcccgcaacgagcgcaaccctcatccttagttgcatcaggtaatgctgagtacttaaggaaactgcc

agtgataagctggaggaaggtggggatgatgtcaagtcatcatggcctttatggagtgggctacacacgtgc

tacaatggtgtctacaatgggttgcaaggtgcgcaagcttaagctaat-ccctaaaagacatctcagttcgg

attgtactctgcaactcgagtacatgaagttggaatcgctagtaatcgtggatcagcatgccacggtgaata

cgttctcgggtct-gtac-----------

>ENA|KJ728742/91-406

ttaagtcccgcaacgagcgcaaccctcatccttagttgcatcaggtaatgctgagtacttaaggaaactgcc

agtgataagctggaggaaggtggggatgatgtcaagtcatcatggcctttatggagtgggctacacacgtgc

tacaatggtgtctacaatgggttgcaaggtgcgcaagcctaagctaat-ccctaaaagacatctcagttcgg

attgtactctgcaactcgagtacatgaagttggaatcgctagtaatcgtggatcatcatgccacggtgaata

cgttctcgggtcttgtacacactgcccgt

>ENA|KJ728743/91-406

ttaagtcccgcaacgagcgcaaccctcatccttagttgcatcaggtaatgctgagtacttaaggaaactgcc

agtgataagctggaggaaggtggggatgatgtcaagtcatcatggcctttatggagtgggctacacacgtgc

tacaatggtgtctacaatgggttgcaaggtgcgcaagcctaagctaat-ccctaaaagacatctcagttcgg

attgtactctgcaactcgagtacatgaagttggaatcgctagtaatcgtggatcagcatgccacggtgaata

cgttctcgggtcttgtacgcactgcccgt

>ENA|KJ728744/91-406

ttaagtcccgcaacgagcgcaaccctcatccttagttgcatcaggtaatgctgagtacttaaggaaactgcc

agtgataagctggaggaaggtggggatgatgtcaagtcatcatggcctttatggagtgggctacacacgtgc

tacaatggtgtctacaatgggttgcaaggtgcgcaagcctaagctaat-ccctaaaagacatctcagttcgg

attgtactctgcaactcgagtacatgaagttggaatcgctagtaatcgtggatcagcatgccacggtgaata

cgttctcgggtcttgtacacactgcccgt

>ENA|KJ728745/91-406

ttaagtcccgcaacgagcgcaaccctcatccttagttacatcaggtaatgctggggacttaaggaaactgcc

agtgataaactggaggaaggtggggatgatgtcaagtcatcatggcccttatggagtgggctacacacgtgc

tacaatggtggctacaatgggctgcaaagt-cgcgaggctaagctaatcccttaaaagccatctcagttcgg

attgtactctgcaactcgagtacatgaagttggaatcgctagtaatcgtggatcagcatgccacggtgaata

cgttctcgggtcttgtacacactgcccgt

>ENA|KJ728748/89-391

ttaagtcccgcaacgagcgcaaccctcatccttagttacatcaggtaatgctggggacttaaggaaactgcc

agtgataaactggaggaaggtggggatgatgtcaagtcatcatggcccttatggagtgggctacacacgtgc

tacaatggtggctacaatgggctgcaaagt-cgcggggctaagctaatcccttaaaagccatctcagttcgg

attgtactctgcaactcgagtgcatgaagttggaatcgctagtaatcgtggatcagcacgccacggtgaata

cg-tctcgggtctgaca------------

>ENA|KJ728750/91-381

ttaagtcccgcaacgagcgcaaccctcatccttagttacatcaggtaatgctggggacttaaggaaactgcc

agtgataaactggaggaaggtggggatgatgtcaagtcatcatggcccttatggagtgggctacacacgtgc

tacaatggtggctacaatgggctgcaaagt-cgcgaggctaagctaatcccttaaaagccatctcagttcgg

attgtactctgcaactcgagtgcatgaagttggaatcgctagtaatcgtggatcagcacgccacggtgaata

cgtt-------------------------

>ENA|KJ728752/91-406

ttaagtcccgcaacgagcgcaaccctcatccttagttacatcaggtaatgctggggacttaaggaaactgcc

agtgataaactggaggaaggtggggatgatgtcaagtcatcatggcccttatggagtgggctacacacgtgc

tacaatggtggctacaatgggctgcaaagt-cgcgaggctaagctaatcccttaaaagccatctcagttcgg

attgtactctgcaactcgagtgcatgaagttggaatcgctagtaatcgtggatcagcacgccacggtgaata

cgttctcgggtcttgtacacactgcccgt

>ENA|KJ728755/50-365

ttaagtcccgcaacgagcgcaaccctcatccttagttacatcaggtaatgctggggacttaaggaaactgcc

agtgataaactggaggaaggtggggatgatgtcaagtcatcatggcccttatggagtgggctacacacgtgc

tacaatggtggctacaatgggctgcaaagt-cgcgaggctaagctaatcccttaaaagccatctcagttcgg

attgtactctgcaactcgagtgcatgaagttggaatcgctagtaatcgtggatcagcacgccacggtgaata

cgttctcgggtcttgtacacactgcccgt

>ENA|KP089991/49-364

ttaagtcccgcaacgagcgcaaccctcatccttagttacatcaggtaatgctggggacttaaggaaactgcc

agtgataaactggaggaaggtggggatgatgtcaagtcatcatggcccttatggagtgggctacacacgtgc

tacaatggtggctacaatgggctgcaaagt-cgcgaggctaagctaatcccttaaaagccatctcagttcgg

attgtactctgcaactcgagtacatgaagttggaatcgctagtaatcgtggatcagcatgccacggtgaata

cgttctcgggtcttgtacacactgcccgt

>ENA|KR698148/70-385

ttaagtcccgcaacgagcgcaaccctcatccttagttgcatcaggtaatgctgagtacttaaggaaactgcc

agtgataagctggaggaaggtggggatgatgtcaagtcatcatggcctttatggagtgggctacacacgtgc

tacaatggtgtctacaatgggctgcaaggtgcgcaagcctaagctaat-ccctaaaagacatctcagttcgg

attgtactctgcaactcgagtacatgaagttggaatcgctagtaatcgtggatcagcatgccacggtgaata

cgttctcgggtcttgtacacactgcccgt

>ENA|KR698151/70-385

ttaagtcccgcaacgagcgcaaccctcatccttagttgcatcaggtaatgctgagtacttaaggaaactgcc

agtgataagctggaggaaggtggggatgatgtcaagtcatcatggcctttatggagtgggctacacacgtgc

tacaatggtgtctacaatgggctgcaaggtgcgcaagcctaagctaat-ccctaaaagacatctcagttcgg

attgtactctgcaactcgagtacatgaagttggaatcgctagtaatcgtggatcagcatgccacggtgaata

cgttctcgggtcttgtacacactgcccgt

>ENA|KR698173/70-385

ttaagtcccgcaacgagcgcaaccctcatccttagttacatcagataatgctggggacttaaggaaactgct

agtgataaactggaggaaggtggggatgatgtcaagtcatcatggcccttatggagtgggctacacacgtgc

tacaatggtggctacaatgggctgcaaagt-cgcgagactaagccaatcccttaaaagccatctcagttcgg

attgtactctgcaactcgagtacatgaagttggaatcgctagtaatcgtggatcagcatgccacggtgaata

cgttctcgggtcttgtacacactgcccgt

>ENA|KT799586/964-1279

ttaagtcccgcaacgagcgcaaccctcatccttagttacatcaggtaatgctggggacttaaggaaactgct

agtgataaactggaggaaggtggggatgatgtcaagtcatcatggcccttatggagtgggctacacacgtgc

tacaatggtggctacaatgggctgcaaagt-cgcgaggctaagctaatcccttaaaagccatctcagttcga

attgcactctgcaactcgagtgcatgaagttggaatcgctagtaatcgtggatcagcatgccacggtgaata

cgttctcgggtcttgtacacactgcccgt

>ENA|KT799588/1008-1323

ttaagtcccgcaacgagcgcaaccctcatccttagttacatcaggtaatgctggggacttaaggaaactgct

agtgataaactggaggaaggtggggatgatgtcaagtcatcatggcccttatggagtgggctacacacgtgc

tacaatggtggctacaataggctgcaaaac-tgcgaagttgagctaatcctttaaaagccatctcagttcgg

attgcactctgcaactcgagtgcatgragttggaatcgctagtaatcgtggatcagcatgccacggtgaata

cgttctcgggtcttgtacacactgcccgt

>ENA|L02888/1037-1352

ttaagtcccgcaacgagcgcaaccctcatccttagttgcatcaggtaatgctgagtacttaaggaaactgcc

agtgataagctggaggaaggtggggatgatgtcaagtcatcatggcctttatggagtgggctacacacgtgc

tacaatggtgtctacaatgggttgcaaggtgcgcaagcctaagctaat-ccctaaaagacatctcagttcgg

attgtactctgcaactcgagtacatgaagttggaatcgctagtaatcgtggatcagcatgccacggtgaata

cgttctcgggtcttgtacacactgcccgt

>ENA|LC108848/1026-1341

ttaagtcccgcaacgagcgcaaccctcatccttagttacatcaggtaatgctggggacttaaggaaactgcc

agtgataaactggaggaaggtggggatgatgtcaagtcatcatggcccttatggagtgggctacacacgtgc

tacaatggtggctacaatgggctgcaaagt-cgcgaggctaagctaatcccttaaaagccatctcagttcgg

attgtactctgcaactcgagtgcatgaagttggaatcgctagtaatcgtggatcagcacgccacggtgaata

cgttctcgggtcttgtacacactgcccgt

>ENA|LC370780/995-1310

ttaagtcccgcaacgagcgcaaccctcatccttagttacatcaggtaatgctggggacttaaggaaactgcc

agtgataaactggaggaaggtggggatgatgtcaagtcatcatggcccttatggagtgggctacacacgtgc

tacaatggtggctacaatgggctgcaaagt-cgcgaggctaagccaatcccttaaaagccatctcagttcgg

attgtactctgcaactcgagtgcatgaagttggaatcgctagtaatcgtggatcagcacgccacggtgaata

cgttctcgggtcttgtacacactgcccgt

>ENA|LC370785/1017-1332

ttaagtcccgcaacgagcgcaaccctcatccttagttacatcaggtaatgctggggacttaaggaaactgcc

agtgataaactggaggaaggtggggatgatgtcaagtcatcatggcccttatggagtgggctacacacgtgc

tacaatggtggctacaatgggctgcaaagt-cgcgaggctaagccaatcccttaaaagccatctcagttcgg

attgtactctgcaactcgagtgcatgaagttggaatcgctagtaatcgtggatcagcacgccacggtgaata

cgttctcgggtcttgtacacactgcccgt

>ENA|MF944114/42-339

ttaagtcccgcaacgagcgcaaccctcatccttagttacatcaggtcatgctggggacttaaggaaactgcc

agtgataaactggaggaaggtggggatgatgtcaagtcatcatggcccttatggagtgggctacacacgtgc

tacaatggtggctacaatgggctgcaaagt-cgcaaggctgagctaat-ccttaaaagccatctcagttcgg

attgtactctgcaactcgagtgcatgaagttggaatcgctagtaatcgtggatcagcacgccacggtgaata

cgttctcgggtc-----------------

>ENA|MF999263/1043-1358

ttaagtcccgcaacgagcgcaaccctcatccttagttgcatcaggtaatgctgagtacttaaggaaactgcc

agtgataagctggaggaaggtggggatgatgtcaagtcatcatggcctttatggagtgggctacacacgtgc

tacaatggtgtctacaatgggttgcaaggtgcgcaagcctaagctaat-ccctaaaagacatctcagttcgg

attgtactctgcaactcgagtacatgaagttggaatcgctagtaatcgtggatcagcatgccacggtgaata

cgttctcgggtcttgtacacactgcccgt

>ENA|MG707893/90-405

ttaagtcccgcaacgagcgcaaccctcatccttagttgcatcaggtaatgctgagtacttaaggaaactgcc

agtgataagctggaggaaggtggggatgatgtcaagtcatcatggcctttatggagtgggctacacacgtgc

tacaatggtgtctacaatgggttgcaaggtgcgcaagcctaagctaat-ccctaaaagacatctcagttcgg

attgtactctgcaactcgagtacatgaagttggaatcgctagtaatcgtggatcagcatgccacggtgaata

cgttctcgggtcttgtacacactgcccgt

>ENA|MG707957/69-382

ttaagtcccgcgacgagcgcaaccctcatccttagttacatcaggttatgctggggacttaaggaaacttcc

agtgataaactggaggaaggtggggatgatgtcaagtcatcacggcccttatggagtgggctacacacgtgc

tacaatggtgattacaatgggctgcaaggt-cgcaaggttgagctaat--cctaaaaatcatctcagttcgg

attgctccccgcaactcgagagcatgaagttggaatcgctagtaatcgtggatcagcgtgccacggtgaata

cgttctcgggtcttgtacacactgcccgt

>ENA|MH470290/537-852

ttaagtcccgcaacgagcgcaaccctcatccttagttgcatcaggtaatgctgagtacttaaggaaactgcc

agtgataagctggaggaaggtggggatgatgtcaagtcatcatggcctttatggagtgggctacacacgtgc

tacaatggtgtctacaatgggctgcaaggtgcgcaagcctaagctaat-ccctaaaagacatctcagttcgg

attgtactctgcaactcgagtacatgaagttggaatcgctagtaatcgtggatcagcatgccacggtgaata

cgttctcgggtcttgtacacactgcccgt

>ENA|MH596695/40-355

ttaagtcccgcaacgagcgcaaccctcatccttagttgcatcaggtaatgctgagcacttaaggaaactgcc

ggtgataagctggaggaagatggggatgatgtcaagtcatcatggcctttatgaagtgggctacacacgtgc

tacaatggtgtctacaatgggctgcaaggtgcgcaagcctaagctaat-ccctaaaagacatctcagttcgg

attgtactctgcaactcgagtgcatgaagttggaatcgctagtaatcgtggatcagcatgccacggtgaata

cgttctcgggtcttgtacacactgcccgt

>ENA|MH596703/40-355

ttaagtcccgcaacgagcgcaaccctcatccttagttgcatcaggtaatgctgagtacttaaggaaactgcc

agtgataagctggaggaaggtggggatgatgtcaagtcatcatggcctttatggagtgggctacacacgtgc

tacaatggtgtctacaatgggctgcaaggtgcgcaagcctaagctaat-ccctaaaagacatctcagttcgg

attgtactctgcaactcgagtacatgaagttggaatcgctagtaatcgtggatcagcatgccacggtgaata

cgttctcgggtcttgtacacactgcccgt

>ENA|MH605279/69-384

ttaagtcccgcaacgagcgcaaccctcatccttagttgcatcaggtaatgctgagtacttaaggaaactgcc

agtgataagctggaggaaggtggggatgatgtcaagtcatcatggcctttatggagtgggctacacacgtgc

tacaatggtgtctacaatgggctgcaaggtgcgcaagcctaagctaat-ccctaaaagacatctcagttcgg

attgtactctgcaactcgagtacatgaagttggaatcgctagtaatcgtggatcagcatgccacggtgaata

cgttctcgggtcttgtacacactgcccgt

>ENA|MK277386/90-405

ttaagtcccgcaacgagcgcaaccctcatccttagttgcatcaggtaatgctgagcacttaaggaaactgcc

agtgataagctggaggaaggtggggatgatgtcaagtcatcatggcctttatgaagtgggctacacacgtgc

tacaatggtgtctacagtgggctgcaaggtgcgcaagcctaagctaat-ccctaaaagacatctcagttcgg

attgtactctgcaactcgagtgcatgaagttggaatcgctagtaatcgtggatcagcatgccacggtgaata

cgttctcgggtcttgtacacactgcccgt

>ENA|MK277388/92-407

ttaagtcccgcaacgagcgcaaccctcatccttagttgcatcaggtaatgctgagcacttaaggaaactgcc

agtgataagctggaggaagatggggatgatgtcaagtcatcatggcctttatgaagtgggctacacacgtgc

tacaatggtgtctacagtgggctgcaaggtgcgcaagcctaagctaat-ccctaaaagacatctcagttcgg

attgtactctgcaactcgagtgcatgaagttggaatcgctagtaatcgtggatcagcatgccacggtgaata

cgttctcgggtcttgtacacactgcccgt

>ENA|MK277393/91-406

ttaagtcccgcaacgagcgcaaccctcatccttagttgcatcaggtaatgctgagtacttaaggaaactgcc

agtgataagctggaggaaggtggggatgatgtcaagtcatcatggcctttatggagtgggctacacacgtgc

tacaatggtgtctacaatgggctgcaaggtgcgcaagcctaagctaat-ccctaaaagacatctcagttcgg

attgtactctgcaactcgagtacatgaagttggaatcgctagtaatcgtggatcagcatgccacggtgaata

cgttctcgggtcttgtacacactgcccgt

>ENA|MK277424/90-405

ttaagtcccgcaacgagcgcaaccctcatccttagttgcatcaggtaatgctgagtacttaaggaaactgcc

agtgataagctggaggaaggtggggatgatgtcaagtcatcatggcctttatggagtgggctacacacgtgc

tacaatggtgtctacaatgggctgcaaggtgcgcaagcctaagctaat-ccctaaaagacatctcagttcgg

attgtactctgcaactcgagtacatgaagttggaatcgctagtaatcgtggatcagcatgccacggtgaata

cgttctcgggtcttgtacacactgcccgt

>ENA|MK278701/70-385

ttaagtcccgcaacgagcgcaaccctcatccttagttgcatcaggtaatgctgagcacttaaggaaactgcc

agtgataagctggaggaagatggggatgatgtcaagtcatcatggcctttatgaagtgggctacacacgtgc

tacaatggtgtctacagtgggctgcaaggtgcgcaagcctaagctaat-ccctaaaagacatctcagttcgg

attgtactctgcaactcgagtgcatgaagttggaatcgctagtaatcgtggatcagcatgccacggtgaata

cgttctcgggtcttgtacacactgcccgt

>ENA|MN123050/984-1299

ttaagtcccgcaacgagcgcaaccctcatccttagttgcatcaggtaatgctgagtacttaaggaaactgcc

agtgataagctggaggaaggtggggatgatgtcaagtcatcatggcctttatggagtgggctacacacgtgc

tacaatggtgtctacaatgggctgcaaggtgcgcaagcctaagctaat-ccctaaaagacatctcagttcgg

attgtactctgcaactcgagtacatgaagttggaatcgctagtaatcgtggatcagcatgccacggtgaata

cgttctcgggtcttgtacacactgcccgt

>ENA|MN268745/84-399

ttaagtcccgcaacgagcgcaaccctcatccttagttgcatcaggtaatgctgagtacttaaggaaactgcc

agtgataagctggaggaaggtggggatgatgtcaagtcatcatggcctttatggagtgggctacacacgtgc

tacaatggtgtctacaatgggctgcaaggtgcgcaagcctaagctaat-ccctaaaagacatctcagttcgg

attgtactctgcaactcgagtacatgaagttggaatcgctagtaatcgtggatcagcatgccacggtgaata

cgttctcgggtcttgtacacactgcccgt

>ENA|MN383058/776-1091

ttaagtcccgcaacgagcgcaaccctcatccttagttgcatcaggtaatgctgagtacttaaggaaactgcc

agtgataagctggaggaaggtggggatgatgtcaagtcatcatggcctttatggagtgggctacacacgtgc

tacaatggtgtctacaatgggctgcaaggtgcgcaagcctaagctaat-ccctaaaagacatctcagttcgg

attgtactctgcaactcgagtacatgaagttggaatcgctagtaatcgtggatcagcatgccacggtgaata

cgttctcgggtcttgtacacactgcccgt

>ENA|MN383129/775-1090

ttaagtcccgcaacgagcgcaaccctcatccttagttgcatcaggtaatgctgagtacttaaggaaactgcc

agtgataagctggaggaaggtggggatgatgtcaagtcatcatggcctttatggagtgggctacacacgtgc

tacaatggtgtctacaatgggctgcaaggtgcgcaagcctaagctaat-ccctaaaagacatctcagttcgg

attgtactctgcaactcgagtacatgaagttggaatcgctagtaatcgtggatcagcatgccacggtgaata

cgttctcgggtcttgtacacactgcccgt

>ENA|MN699328/957-1272

ttaagtcccgcaacgagcgcaaccctcatccttagttacatcaggtaatgctggggacttaaggaaactgct

agtgataaactggaggaaggtggggatgatgtcaagtcatcatggcccttatggagtgggctacacacgtgc

tacaatggtggctacaatgggctgcaaagt-cgcgaggctaagctaatcccttaaaagccatctcagttcga

attgcactctgcaactcgagtgcatgaagttggaatcgctagtaatcgtggatcagcatgccacggtgaata

cgttctcgggtcttgtacacactgcccgt

>ENA|MN928707/90-405

ttaagtcccgcaacgagcgcaaccctcatccttagttgcatcaggtaatgctgagcacttaaggaaactgcc

agtgataagctggaggaagatggggatgatgtcaagtcatcatggcctttatgaagtgggctacacacgtgc

tacaatggtgtctacagtgggctgcaaggtgcgcaagcctaagctaat-ccctaaaagacatctcagttcgg

attgtactctgcaactcgagtgcatgaagttggaatcgctagtaatcgtggatcagcatgccacggtgaata

cgttctcgggtcttgtacacactgcccgt

>ENA|MW179590/69-384

ttaagtcccgcaacgagcgcaaccctcatccttagttgcatcaggtaatgctgagtacttaaggaaactgcc

agtgataagctggaggaaggtggggatgatgtcaagtcatcatggcctttatggagtgggctacacacgtgc

tacaatggtgtctacaatgggttgcaaggtgcgcaagcttaagctaat-ccctaaaagacatctcagttcgg

attgtactctgcaactcgagtacatgaagttggaatcgctagtaatcgtggatcagcatgccacggtgaata

cgttctcgggtcttgtacacactgcccgt

>ENA|MZ031918/92-407

ttaagtcccgcaacgagcgcaaccctcatccttagttgcatcaggtaatgctgagcacttaaggaaactgcc

agtgataagctggaggaagatggggatgatgtcaagtcatcatggcctttatgaagtgggctacacacgtgc

tacaatggtgtctacaatgggctgcaaggtgcgcaagcctaagctaat-ccctaaaagacatctcagttcgg

attgtactctgcaactcgagtgcatgaagttggaatcgctagtaatcgtggatcagcatgccacggtgaata

cgttctcgggtcttgtacacactgcccgt

>ENA|MZ031920/92-407

ttaagtcccgcaacgagcgcaaccctcatccttagttgcatcaggtaatgctgagcacttaaggaaactgcc

agtgataagctggaggaagatggggatgatgtcaagtcatcatggcctttatgaagtgggctacacacgtgc

tacaatggtgtctacaatgggctgcaaggtgcgcaagcctaagctaat-ccctaaaagacatctcagttcgg

attgtactctgcaactcgagtgcatgaagttggaatcgctagtaatcgtggatcagcatgccacggtgaata

cgttctcgggtcttgtacacactgcccgt

>CP031221:1173014-1174384/998-1313

ttaagtcccgcaacgagcgcaaccctcatccttagttgcatcaggtaatgctgagtacttaaggaaactgcc

agtgataagctggaggaaggtggggatgatgtcaagtcatcatggcctttatggagtgggctacacacgtgc

tacaatggtgtctacaatgggttgcaaggtgcgcaagcctaagctaat-ccctaaaagacatctcagttcgg

attgtactctgcaactcgagtacatgaagttggaatcgctagtaatcgtggatcagcatgccacggtgaata

cgttctcgggtcttgtacacactgcccgt

>CP021120.1:596706-598076/998-1313

ttaagtcccgcaacgagcgcaaccctcatccttagttgcatcaggtaatgctgagtacttaaggaaactgcc

agtgataagctggaggaaggtggggatgatgtcaagtcatcatggcctttatggagtgggctacacacgtgc

tacaatggtgtctacaatgggctgcaaggtgcgcaagcctaagctaat-ccctaaaagacatctcagttcgg

attgtactctgcaactcgagtacatgaagttggaatcgctagtaatcgtggatcagcatgccacggtgaata

cgttctcgggtcttgtacacactgcccgt

>SAWADOGO_4-16S/49-364

ttaagtcccgcaacgagcgcaaccctcatccttagttgcatcaggtaatgctgagtacttaaggaaactgcc

agtgataagctggaggaaggtggggatgatgtcaagtcatcatggcctttatggagtgggctacacacgtgc

tacaatggtgtctacaatgggttgcaaggtgcgcaagcctaagctaat-ccctaaaagacatctcagttcgg

attgtactctgcaactcgagtacatgaagttggaatcgctagtaatcgtggatcagcatgccacggtgaata

cgttctcgggtcttgtacacactgcccgt

>SAWADOGO_5-16S/47-362

ttaagtcccgcaacgagcgcaaccctcatccttagttgcatcaggtaatgctgagtacttaaggaaactgcc

agtgataagctggaggaaggtggggatgatgtcaagtcatcatggcctttatggagtgggctacacacgtgc

tacaatggtgtctacaatgggttgcaaggtgcgcaagcctaagctaat-ccctaaaagacatctcagttcgg

attgtactctgcaactcgagtacatgaagttggaatcgctagtaatcgtggatcagcatgccacggtgaata

cgttctcgggtcttgtacacactgcccgt

>SAWADOGO_6-16S/49-364

ttaagtcccgcaacgagcgcaaccctcatccttagttgcatcaggtaatgctgagtacttaaggaaactgcc

agtgataagctggaggaaggtggggatgatgtcaagtcatcatggcctttatggagtgggctacacacgtgc

tacaatggtgtctacaatgggttgcaaggtgcgcaagcctaagctaat-ccctaaaagacatctcagttcgg

attgtactctgcaactcgagtacatgaagttggaatcgctagtaatcgtggatcagcatgccacggtgaata

cgttctcgggtcttgtacacactgcccgt

>SAWADOGO_8-16S/46-361

ttaagtcccgcaacgagcgcaaccctcatccttagttgcatcaggtaatgctgagtacttaaggaaactgcc

agtgataagctggaggaaggtggggatgatgtcaagtcatcatggcctttatggagtgggctacacacgtgc

tacaatggtgtctacaatgggctgcaaggtgcgcaagcctaagctaat-ccctaaaagacatctcagttcgg

attgtactctgcaactcgagtacatgaagttggaatcgctagtaatcgtggatcagcatgccacggtgaata

cgttctcgggtcttgtacacactgcccgt

>SAWADOGO_9-16S/47-362

ttaagtcccgcaacgagcgcaaccctcatccttagttgcatcaggtaatgctgagtacttaaggaaactgcc

agtgataagctggaggaaggtggggatgatgtcaagtcatcatggcctttatggagtgggctacacacgtgc

tacaatggtgtctacaatgggctgcaaggtgcgcaagcctaagctaat-ccctaaaagacatctcagttcgg

attgtactctgcaactcgagtacatgaagttggaatcgctagtaatcgtggatcagcatgccacggtgaata

cgttctcgggtcttgtacacactgcccgt

>SAWADOGO_10-16S/48-363

ttaagtcccgcaacgagcgcaaccctcatccttagttgcatcaggtaatgctgagtacttaaggaaactgcc

agtgataagctggaggaaggtggggatgatgtcaagtcatcatggcctttatggagtgggctacacacgtgc

tacaatggtgtctacaatgggctgcaaggtgcgcaagcctaagctaat-ccctaaaagacatctcagttcgg

attgtactctgcaactcgagtacatgaagttggaatcgctagtaatcgtggatcagcatgccacggtgaata

cgttctcgggtcttgtacacactgcccgt

>SAWADOGO_12-16S/48-363

ttaagtcccgcaacgagcgcaaccctcatccttagttgcatcaggtaatgctgagtacttaaggaaactgcc

agtgataagctggaggaaggtggggatgatgtcaagtcatcatggcctttatggagtgggctacacacgtgc

tacaatggtgtctacaatgggctgcaaggtgcgcaagcctaagctaat-ccctaaaagacatctcagttcgg

attgtactctgcaactcgagtacatgaagttggaatcgctagtaatcgtggatcagcatgccacggtgaata

cgttctcgggtcttgtacacactgcccgt

>SAWADOGO_13-16S/48-363

ttaagtcccgcaacgagcgcaaccctcatccttagttgcatcaggtaatgctgagtacttaaggaaactgcc

agtgataagctggaggaaggtggggatgatgtcaagtcatcatggcctttatggagtgggctacacacgtgc

tacaatggtgtctacaatgggttgcaaggtgcgcaagcctaagctaat-ccctaaaagacatctcagttcgg

attgtactctgcaactcgagtacatgaagttggaatcgctagtaatcgtggatcagcatgccacggtgaata

cgttctcgggtcttgtacacactgcccgt

>SAWADOGO_15-16S/48-363

ttaagtcccgcaacgagcgcaaccctcatccttagttgcatcaggtaatgctgagcacttaaggaaactgcc

agtgataagctggaggaagatggggatgatgtcaagtcatcatggcctttatgaagtgggctacacacgtgc

tacaatggtgtctacaatgggctgcaaggtgcgcaagcctaagctaat-ccctaaaagacgtctcagttcgg

attgtactctgcaactcgagtgcatgaagttggaatcgctagtaatcgtggatcagcatgccacggtgaata

cgttctcgggtcttgtacacactgcccgt

>SAWADOGO_16-16S/47-362

ttaagtcccgcaacgagcgcaaccctcatccttagttgcatcaggtaatgctgagtacttaaggaaactgcc

agtgataagctggaggaaggtggggatgatgtcaagtcatcatggcctttatggagtgggctacacacgtgc

tacaatggtgtctacaatgggttgcaaggtgcgcaagcctaagctaat-ccctaaaagacatctcagttcgg

attgtactctgcaactcgagtacatgaagttggaatcgctagtaatcgtggatcagcatgccacggtgaata

cgttctcgggtcttgtacacactgcccgt

>SAWADOGO_18-16S/48-363

ttaagtcccgcaacgagcgcaaccctcatccttagttgcatcaggtaatgctgagtacttaaggaaactgcc

agtgataagctggaggaaggtggggatgatgtcaagtcatcatggcctttatggagtgggctacacacgtgc

tacaatggtgtctacaatgggttgcaaggtgcgcaagcctaagctaat-ccctaaaagacatctcagttcgg

attgtactctgcaactcgagtacatgaagttggaatcgctagtaatcgtggatcagcatgccacggtgaata

cgttctcgggtcttgtacacactgcccgt

>SAWADOGO_21-16S/48-363

ttaagtcccgcaacgagcgcaaccctcatccttagttgcatcaggtaatgctgagtacttaaggaaactgcc

agtgataagctggaggaaggtggggatgatgtcaagtcatcatggcctttatggagtgggctacacacgtgc

tacaatggtgtctacaatgggttgcaaggtgcgcaagcctaagctaat-ccctaaaagacatctcagttcgg

attgtactctgcaactcgagtacatgaagttggaatcgctagtaatcgtggatcagcatgccacggtgaata

cgttctcgggtcttgtacacactgcccgt

>SAWADOGO_23-16S/48-363

ttaagtcccgcaacgagcgcaaccctcatccttagttgcatcaggtaatgctgagtacttaaggaaactgcc

agtgataagctggaggaaggtggggatgatgtcaagtcatcatggcctttatggagtgggctacacacgtgc

tacaatggtgtctacaatgggttgcaaggtgcgcaagcctaagctaat-ccctaaaagacatctcagttcgg

attgtactctgcaactcgagtacatgaagttggaatcgctagtaatcgtggatcagcatgccacggtgaata

cgttctcgggtcttgtacacactgcccgt

>SAWADOGO_24-16S/47-362

ttaagtcccgcaacgagcgcaaccctcatccttagttgcatcaggtaatgctgagtacttaaggaaactgcc

agtgataagctggaggaaggtggggatgatgtcaagtcatcatggcctttatggagtgggctacacacgtgc

tacaatggtgtctacaatgggttgcaaggtgcgcaagcctaagctaat-ccctaaaagacatctcagttcgg

attgtactctgcaactcgagtacatgaagttggaatcgctagtaatcgtggatcagcatgccacggtgaata

cgttctcgggtcttgtacacactgcccgt

>SAWADOGO_25-16S/48-363

ttaagtcccgcaacgagcgcaaccctcatccttagttgcatcaggtaatgctgagtacttaaggaaactgcc

agtgataagctggaggaaggtggggatgatgtcaagtcatcatggcctttatggagtgggctacacacgtgc

tacaatggtgtctacaatgggttgcaaggtgcgcaagcctaagctaat-ccctaaaagacatctcagttcgg

attgtactctgcaactcgagtacatgaagttggaatcgctagtaatcgtggatcagcatgccacggtgaata

cgttctcgggtcttgtacacactgcccgt

>SAWADOGO_26-16S/48-363

ttaagtcccgcaacgagcgcaaccctcatccttagttgcatcaggtaatgctgagtacttaaggaaactgcc

agtgataagctggaggaaggtggggatgatgtcaagtcatcatggcctttatggagtgggctacacacgtgc

tacaatggtgtctacaatgggttgcaaggtgcgcaagcctaagctaat-ccctaaaagacatctcagttcgg

attgtactctgcaactcgagtacatgaagttggaatcgctagtaatcgtggatcagcatgccacggtgaata

cgttctcgggtcttgtacacactgcccgt

>SAWADOGO_28-16S/49-364

ttaagtcccgcaacgagcgcaaccctcatccttagttgcatcaggtaatgctgagtacttaaggaaactgcc

agtgataagctggaggaaggtggggatgatgtcaagtcatcatggcctttatggagtgggctacacacgtgc

tacaatggtgtctacaatgggttgcaaggtgcgcaagcctaagctaat-ccctaaaagacatctcagttcgg

attgtactctgcaactcgagtacatgaagttggaatcgctagtaatcgtggatcagcatgccacggtgaata

cgttctcgggtcttgtacacactgcccgt

>SAWADOGO_C+-16S/48-363

ttaagtcccgcaacgagcgcaaccctcatccttagttgcatcaggtaatgctgagtacttaaggaaactgcc

agtgataagctggaggaaggtggggatgatgtcaagtcatcatggcctttatggagtgggctacacacgtgc

tacaatggtgtctacaatgggctgcaaggtgcgcaagcctaagctaat-ccctaaaagacatctcagttcgg

attgtactctgcaactcgagtacatgaagttggaatcgctagtaatcgtggatcagcatgccacggtgaata

cgttctcgggtcttgtacacactgcccgt
